# Mesenchymal Lineage Heterogeneity Underlies Non-Redundant Functions of Pancreatic Cancer-Associated Fibroblasts

**DOI:** 10.1101/2021.05.01.442252

**Authors:** Erin J. Helms, Mark W. Berry, R. Crystal Chaw, Chris C. DuFort, Duanchen Sun, M. Kathrina Onate, Chet Oon, Sohinee Bhattacharyya, Hannah Sanford-Crane, Wesley Horton, Jennifer M. Finan, Ariana Sattler, Rosemary Makar, David W. Dawson, Zheng Xia, Sunil R. Hingorani, Mara H. Sherman

## Abstract

Cancer-associated fibroblast (CAF) heterogeneity is increasingly appreciated, but the origins and functions of distinct CAF subtypes remain poorly understood. The abundant and transcriptionally diverse CAF population in pancreatic ductal adenocarcinoma (PDAC) is thought to arise from a common cell of origin, pancreatic stellate cells (PSCs), with diversification resulting from cytokine and growth factor gradients within the tumor microenvironment. Here we analyzed the differentiation and function of PSCs during tumor progression *in vivo*. Contrary to expectations, we found that PSCs give rise to a numerically minor subset of PDAC CAFs. Targeted ablation of PSC-derived CAFs within their host tissue revealed non-redundant functions for this defined CAF population in shaping the PDAC microenvironment, including production of specific components of the extracellular matrix. Together, these findings link stromal evolution from distinct cells of origin to transcriptional heterogeneity among PDAC CAFs, and demonstrate unique functions for CAFs of a defined cellular origin.

**Statement of significance:** By tracking and ablating a specific CAF population, we find that a numerically minor CAF subtype from a defined cell of origin plays unique roles in establishing the pancreatic tumor microenvironment. Together with prior studies, this work suggests that mesenchymal lineage heterogeneity as well as signaling gradients diversify PDAC CAFs.

## Introduction

Pancreatic ductal adenocarcinoma (PDAC) is defined in part by an exuberant stromal reaction, including abundant cancer-associated fibroblasts (CAFs) (1-4). Diverse tumor-supportive functions have been ascribed to PDAC CAFs, including metabolic roles whereby nutrient transfer from CAFs to neighboring pancreatic cancer cells facilitates proliferation within a nutrient-poor microenvironment (5-9). In addition, PDAC CAFs produce cytokines and chemokines associated with immune suppression (10-13), and CAF ablation in mice fosters efficacy of immune checkpoint inhibitors which are otherwise ineffective in PDAC (14, 15). Further, perhaps the best known function of activated fibroblasts in a wound-healing reaction and of CAFs in a tumor microenvironment is to produce extracellular matrix (ECM) components and remodeling enzymes. The dense ECM in PDAC physically impedes the vasculature and limits delivery of intravenous therapeutic agents (16, 17), and PDAC patients with high levels of stiff fibrosis enriched for matricellular proteins (such as tenascin C) have shortened survival (18). These findings have motivated efforts to develop therapies targeting CAFs, including inhibitors of pathways that regulate their phenotypes (e.g., the Hedgehog pathway) (19, 20) or their immune-modulatory functions (e.g., the CXCL12-CXCR4 axis) (14, 21), and agents targeting the ECM itself (22).

However, the impact of CAFs on PDAC progression and the therapeutic potential of targeting these cells are controversial. Genetic or pharmacologic ablation of CAFs during PDAC progression in mice—either targeting alpha-smooth muscle actin (α-SMA)-expressing CAFs (15) or Sonic Hedgehog-dependent CAFs (23, 24)—resulted in poorly differentiated tumors, and caused mice to succumb to disease faster than those with CAF-replete PDAC. In addition, stromal depletion of ECM component type I collagen significantly accelerated mortality in PDAC-bearing mice (25), and inhibition of collagen crosslinking by LOXL2 increased PDAC growth and reduced overall survival (26). Similarly, higher tumor stromal density, including cellular and acellular components of the stroma, associated with longer overall survival among PDAC patients (26, 27). These studies highlight the tumor-suppressive or homeostatic potential of PDAC CAFs, and impel a more thorough understanding of this complex compartment of the tumor microenvironment with respect to PDAC progression.

To reconcile these seemingly contradictory findings, several groups have postulated that PDAC CAFs are heterogeneous, potentially including subtypes that support and others that suppress tumor growth. Consistent with this notion, single-cell RNA-seq and other approaches have revealed transcriptional heterogeneity among CAFs in murine and human PDAC (11, 12, 28-30). Important considerations moving forward are the origins of these CAF subtypes and, importantly, their functions. Understanding the potentially unique functions of CAF subtypes is important in hopes of identifying and specifically targeting the tumor-promoting mechanisms in the stroma. Understanding CAF cellular origin(s) is important because blocking the development or activation or tumor-supportive CAF subtypes may be a viable therapeutic strategy. Identifying CAF origin is also important for the goal of developing new models to track and manipulate CAFs within their host tissue in a robust and specific way, as such models are presently lacking. PDAC CAFs are generally thought to share a common cell of origin: a tissue-resident mesenchymal cell called a stellate cell (31, 32). Stellate cells are found in two tissues in the body, the liver and the pancreas (33); while two Cre-based models have informed on hepatic stellate cell fate and function, these models have not been used successfully to study pancreatic stellate cells (PSCs). As a result, our understanding of PSC biology to date has been extrapolated from the liver or learned from cell culture studies.

In healthy pancreas tissue, PSCs are in a quiescent state, characterized in part by cytoplasmic lipid droplets that store vitamin A as retinyl esters, and implicated in tissue homeostasis including recycling of the basement membrane (34). Upon exposure to tissue damage cues or a stiff growth substrate, PSCs become activated, and trans-differentiated to a myofibroblastic phenotype. As PSCs can be isolated from healthy pancreata by density centrifugation on the basis of their lipid content, PSC activation can be modeled *in vitro*. Transcriptional profiling of PSC activation showed upregulation of ECM components and remodeling enzymes, growth factors, and other signatures associated with CAFs, suggesting that PSCs are indeed competent to give rise to CAFs during PDAC progression. However, as these cells have not been tracked *in vivo* in the context of tumorigenesis, their contribution to the PDAC microenvironment and their functions therein remain unknown. Recent work demonstrated that PDAC CAF heterogeneity results in part from signaling gradients of critical factors including TGF-β and IL-1 (12, 35), and that these factors can differentially program PSCs into inflammatory or myofibroblastic CAF fates *in vitro*. To analyze the fate of PSCs in PDAC *in vivo*, we developed a mouse model in which we can track and specifically ablate these cells within the pancreas, leveraging their unique lipid-storing origin. We find that PSCs indeed give rise to CAFs, but that this PSC-derived CAF population is numerically minor, suggesting additional and yet-undefined cellular origins for the majority of PDAC CAFs. Importantly, PSC-derived CAFs play significant, non-redundant roles in modulation of the tumor microenvironment including production of specific components of the ECM, suggesting that these cells or their critical regulators may be therapeutic targets.

## Results

### Characterization of a genetic approach to label and track stellate cells in normal pancreas tissue

To identify a Cre-based approach to label and track PSCs *in vivo*, we analyzed published RNA-seq data from primary PSCs and noted very high expression of adipocyte marker fatty acid binding protein 4 (Fabp4) in the quiescent state (36). While Fabp4 expression was dramatically downregulated in the activated, fibroblastic state, we reasoned that a lineage labeling approach driven by Fabp4 regulatory elements would label PSCs as well as CAFs derived from them. To address the utility of this approach, we crossed *Fabp4-Cre* mice, developed to study adipose tissue (37), to mice harboring the *Rosa26*^*ACTB-tdTomato,-EGFP*^ (*Rosa26*^*mTmG*^ hereafter) reporter allele (38) (Fig. 1A). In these mice, the ubiquitous *Rosa26* promoter drives expression of tdTomato followed by a stop codon; in the presence of Cre, tdTomato and the stop codon are excised, leading to expression of GFP. As such, all cells of the mouse express tdTomato except Cre-positive cells and their progeny, which are indelibly labeled with GFP. When we examined pancreata of *Fabp4-Cre;Rosa*^*mTmG*^ mice, we noted rare GFP^+^ cells in the periacinar spaces of the expected morphology and frequency for PSCs based on previously published PSC characterization and electron microscopy analyses (39, 40) (Fig. 1B). We characterized these GFP^+^ cells in the pancreas to assess whether this lineage label was specific to PSCs within normal pancreas tissue, and whether GFP labeling was pervasive among PSCs. To assess specificity, we analyzed GFP together with markers of other known pancreatic cell types, as strong markers for PSCs are largely lacking. We found that CD31^+^ endothelial cells, CD45^+^ leukocytes, and NG2^+^ pericytes were restricted to the tdTomato^+^ population and lacked the GFP lineage label (Fig. 1C-E, Supplementary Fig. S1A). Though CD31^+^ endothelial cells were all tdTomato^+^, we noted a perivascular localization of a subset of PSCs, as has been reported for HSCs (Fig. 1C) (41). To further address specificity, we isolated tdTomato^+^ and GFP^+^ cells from the pancreas by FACS (Supplementary Fig. 1B) and measured expression of markers for pancreatic cell types by qPCR (Fig. 1F). We did not see an enrichment for *Cspg4* (which encodes pericyte marker NG2) in the GFP^+^ fraction, suggesting that the numerically small pericyte population is within the far more numerous tdTomato^+^ cell population (liver was a positive control). Markers of acinar cells, ductal cells, and beta cells were restricted to the tdTomato^+^ fraction, while mesenchymal marker *Vim* and putative stellate cell/mesenchymal marker *Des* were strongly enriched in the GFP^+^ fraction. These results suggest that GFP specifically marks PSCs within the pancreas of *Fabp4-Cre;Rosa26*^*mTmG*^ mice.

**Figure 1.**
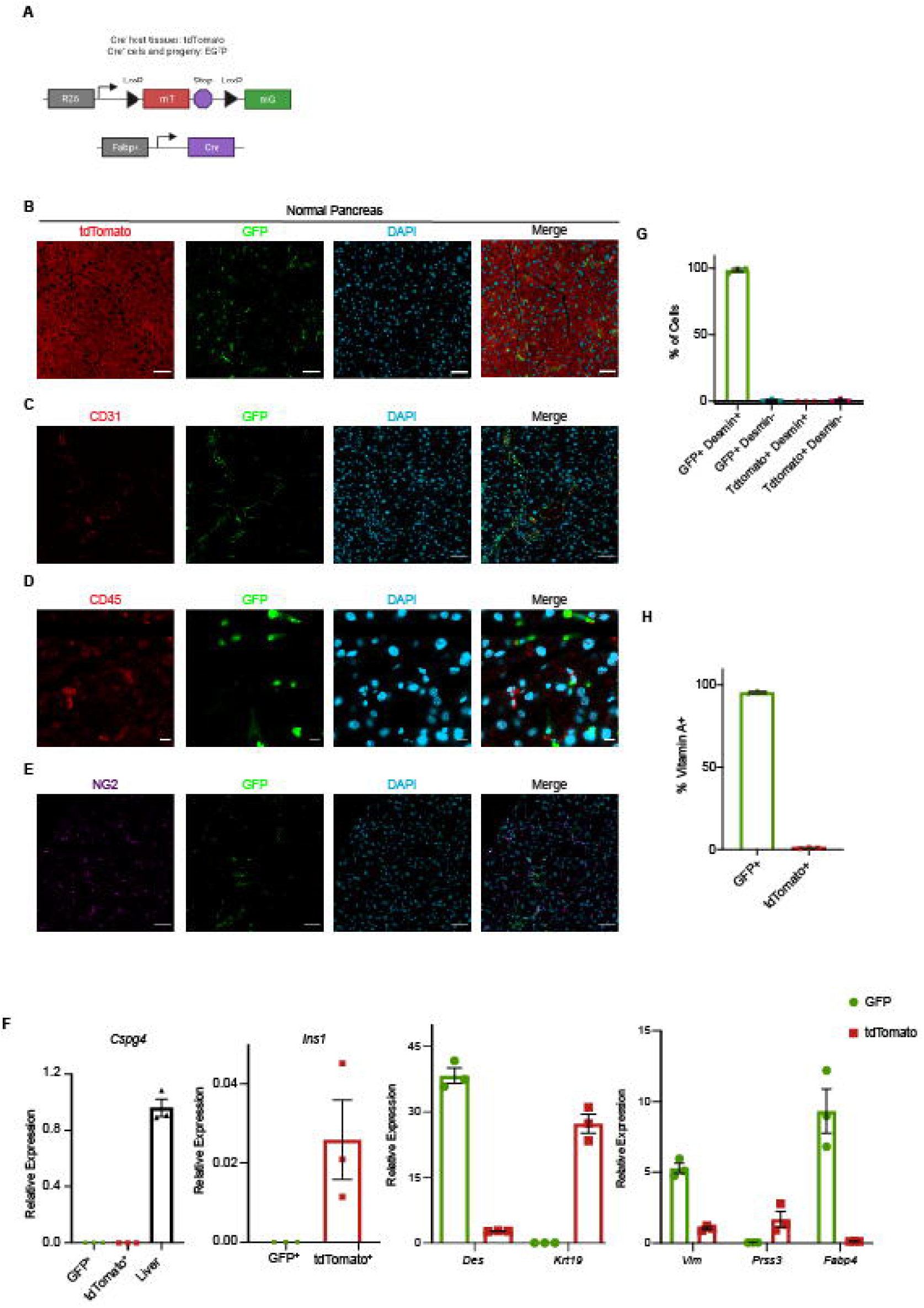
Fabp4-Cre marks stellate cells specifically and pervasively within normal pancreas tissue. **(A**) Schematic of the alleles used to label and track PSCs *in vivo*. **(B)** Representative image of normal pancreas tissue from *Fabp4-Cre;Rosa26*^*mTmG*^ mice (n = 7) showing rare GFP^+^ cells within a predominantly tdTomato^+^ tissue. Scale bar = 50 μm. **(C)** Representative image of normal pancreas tissue from *Fabp4-Cre;Rosa26*^*mTmG*^ mice (n = 3) showing CD31^+^ endothelial cells and GFP^+^ cells, some of which are adjacent to vessels. Scale bar = 50 μm. **(D)** Representative image of normal pancreas tissue from *Fabp4-Cre;Rosa26*^*mTmG*^ mice (n = 3) showing CD45^+^ leukocytes and GFP^+^ cells. Scale bar = 10 μm. **(E)** Representative image of normal pancreas tissue from *Fabp4-Cre;Rosa26*^*mTmG*^ mice (n = 3) showing NG2^+^ pericytes and GFP^+^ cells. Scale bar = 50 μm. **(F)** qPCR for the indicated genes in GFP^+^ and tdTomato^+^ cells isolated from normal pancreas tissue from *Fabp4-Cre;Rosa26*^*mTmG*^ mice by FACS (n = 3, with each replicate pooled from 2 mice), including markers of pericytes (*Cspg4*; liver is a positive control), stellate cells and potentially other mesenchymal cells (*Des*), ductal cells (*Krt19*), mesenchymal cells (*Vim*), acinar cells (*Prss3*), and beta cells (*Ins1*); *Fabp4* was included as a control. Data were normalized to *36b4* and are presented as mean ± SEM. **(G)** Quantification of GFP^+^ cells and tdTomato^+^ cells out of total, Desmin^+^ PSCs isolated by density centrifugation from normal pancreas tissue in *Fabp4-Cre;Rosa26*^*mTmG*^ mice (n = 3) and analyzed by immunofluorescence microscopy. Data are presented as mean ± SEM. **(H)** Flow cytometry results depicting GFP^+^ and tdTomato^+^ cells among all vitamin A^+^ PSCs in normal pancreas tissue from *Fabp4-Cre;Rosa26*^*mTmG*^ mice (n = 3). Data are presented as mean ± SEM.

We next assessed whether GFP pervasively marks PSCs in *Fabp4-Cre;Rosa26*^*mTmG*^ mice, which we analyzed in two ways. We isolated PSCs from the pancreas by density centrifugation, which broadly captures quiescent PSCs on the basis of their lipid content. As this is an enrichment but not a purification, we plated cells at the interface and stained for Desmin, to increase confidence that our analysis extended to all PSCs but not to any contaminating cell types of similar density. We found that nearly all PSCs identified by this approach were GFP^+^ (Fig. 1G, Supplementary Fig. S1C). To analyze this a second way, we noted that perhaps the best-known function of quiescent stellate cells is storage of vitamin A in their cytoplasmic lipid droplets as retinyl esters. This retinoid storage gives stellate cells a blue-green autofluorescence that can be analyzed by flow cytometry, as previously demonstrated for hepatic stellate cells (42). We observed the expected frequency of vitamin A^+^ PSCs upon flow cytometric analysis of a single cell suspension of normal pancreas from *Fabp4-Cre;Rosa26*^*mTmG*^ mice (Supplementary Fig. 1D). When we analyzed our lineage labels among these vitamin A^+^ PSCs, we found that nearly all were GFP^+^ (Fig. 1H). These results together suggest that *Fabp4-Cre;Rosa26*^*mTmG*^ mice feature specific and pervasive GFP labeling of stellate cells within the pancreas.

### Analysis of stellate cell contribution to the CAF pool in the PDAC microenvironment

As our model allows us to track the fate of PSCs during pancreatic tumorigenesis, we used *Fabp4-Cre;Rosa26*^*mTmG*^ mice to formally address the contribution of PSCs to the PDAC CAF population. Upon transplantation of *Kras*^*LSL-G12D/+*^*;Trp53*^*LSL-R172H/+*^*;Pdx1-Cre* (KPC) PDAC cells into the pancreas of *Fabp4-Cre;Rosa26*^*mTmG*^ hosts, we consistently observed an expansion of GFP^+^ cells in the tumor microenvironment with a CAF-like morphology (Fig. 2A). To characterize these GFP^+^ stromal cells, we stained for CAF markers including pan-CAF marker Podoplanin (PDPN) (11) and myofibroblastic CAF marker α-SMA (28). We found that GFP^+^ or PSC-derived CAFs expressed these markers (Fig. 2B,C), confirming that PSCs give rise to PDAC CAFs and demonstrating that they yield a subset of CAFs within the previously established myCAF subpopulation. Contrary to our expectations, however, we noted across PDAC models that PSCs give rise to only a small minority of CAFs, raising the possibility that previously described transcriptional heterogeneity among these cells is due in part to distinct cells of origin. We confirmed these findings by flow cytometry, which showed that, in two different KPC-derived models and using two different CAF cell surface markers (PDPN or PDGFRα), PSC-derived CAFs give rise to approximately 10-15% of total PDPN^+^ CAFs (Fig. 2D-F). These results indicate that PSCs give rise to a numerically minor subset of PDAC CAFs, and prompted us to examine whether this PSC-derived CAF population plays unique roles in the tumor microenvironment, or whether these CAFs harbor similar transcriptional profiles and functions to CAFs of other cellular origins.

**Figure 2.**
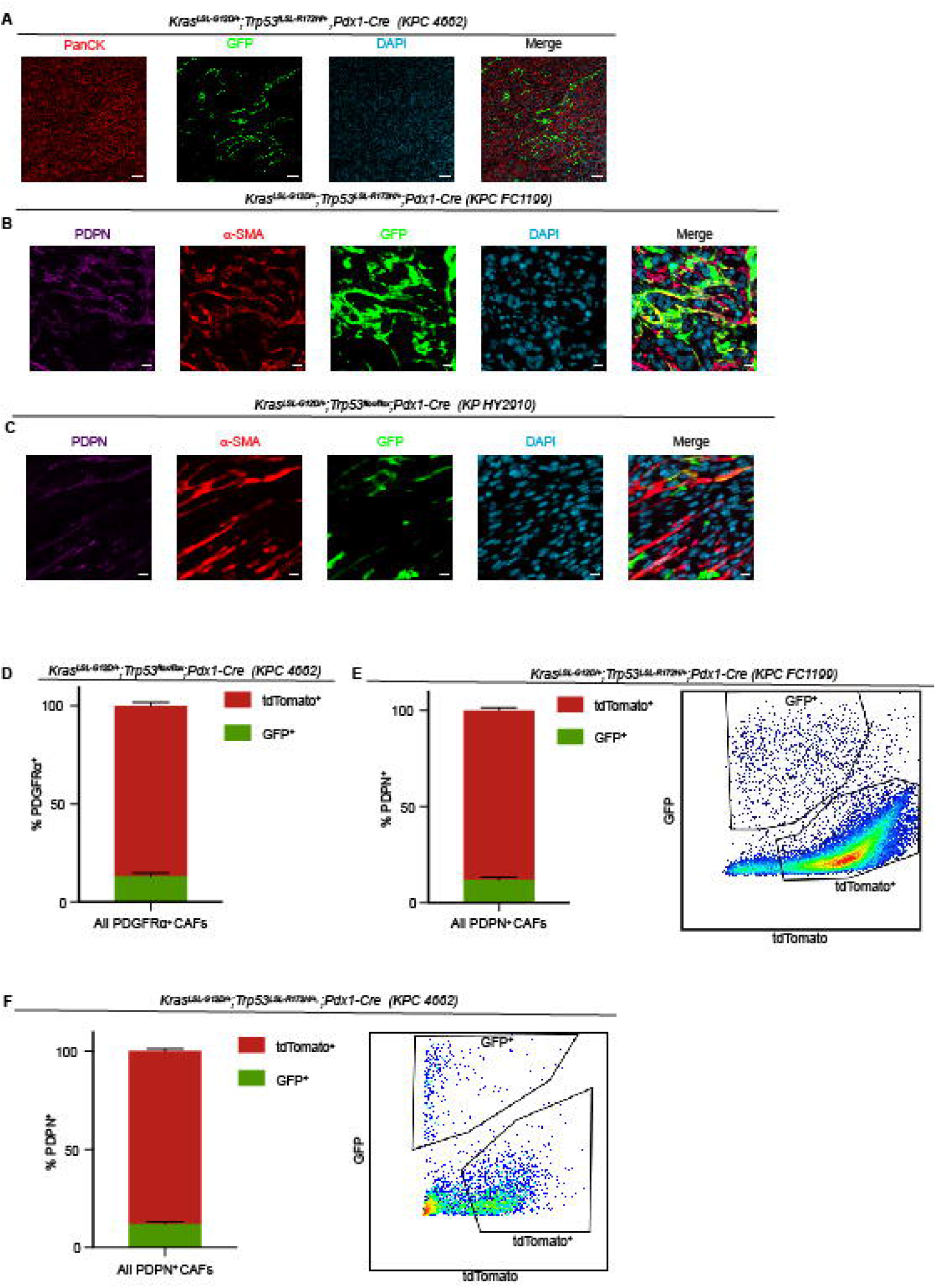
Stellate cells give rise to a numerically minor subset of PDAC CAFs. **(A)** Immunohistochemical staining of PDAC (KPC 4662) in *Fabp4-Cre;Rosa26*^*mTmG*^ hosts (n = 5), with GFP in green and panCK (tumor cells) in red. Scale bar = 50 μm. **(B)** Immunohistochemical staining of PDAC (KPC FC1199) in *Fabp4-Cre;Rosa26*^*mTmG*^ hosts (n = 3), stained for GFP, PDPN, and α-SMA. Scale bar = 10 μm. **(C)** Immunohistochemical staining of PDAC (KP^flox/+^C HY2910) in *Fabp4-Cre;Rosa26*^*mTmG*^ hosts (n = 3), stained for GFP, PDPN, and α-SMA. Scale bar = 10 μm. **(D)** Flow cytometry analysis of PDGFRα, GFP, and tdTomato in KPC 4662 tumors in *Fabp4-Cre;Rosa26*^*mTmG*^ hosts (n = 5). Data are presented as mean ± SEM. **(E)** Flow cytometry analysis of PDPN, GFP, and tdTomato in KPC FC1199 tumors in *Fabp4-Cre;Rosa26*^*mTmG*^ hosts (n = 8). Data are presented as mean ± SEM. **(F)** Flow cytometry analysis of PDPN, GFP, and tdTomato in KPC 4662 tumors in *Fabp4-Cre;Rosa26*^*mTmG*^ hosts (n = 3). Data are presented as mean ± SEM.

### Distinct transcriptional profiles of CAFs from a stellate cell or non-stellate-cell origin

To assess the potential non-redundancy of PSC-derived CAFs, we analyzed their transcriptional profiles. To this end, we established PDAC in *Fabp4-Cre;Rosa26*^*mTmG*^ hosts, isolated GFP^+^ (PSC-derived) and tdTomato^+^ (non-PSC-derived) CAFs by FACS, and analyzed gene expression by RNA-seq. This analysis revealed that both CAF populations express similar levels of broad or pan-CAF markers including *Fap* (which encodes fibroblast activation protein) and *Pdpn*, and that both populations expressed very high levels of *Acta2* (which encodes α-SMA). This transcriptional similarity extended to the majority of collagen genes. However, we found extensive, significant transcriptional differences between these CAF populations (Fig. 3A), suggesting that transcriptional heterogeneity among PDAC CAFs is not only a consequence of cytokine and growth factor gradients within the tumor microenvironment, but also of mesenchymal lineage heterogeneity. Gene ontology analysis revealed that genes more highly expressed in the PSC-derived CAF population are significantly enriched for those involved in cell adhesion, ECM-receptor interaction, and axon guidance (Fig. 3B). The gene identities on these lists included cell surface adhesion molecules that facilitate leukocyte trafficking and/or cancer cell spatial patterning but have yet to be characterized on CAFs, including the receptor tyrosine kinase *Tie1* (Fig. 3C) (notably, *Tek* encoding TIE2 was not enriched among PSC-derived CAFs). ECM components more highly or uniquely expressed among PSC-derived CAFs include those implicated in tissue stiffness and PDAC aggressiveness, such as tenascins including *Tnc* (Fig. 3C) (18) and *Hspg2*, encoding perlecan. Differential expression of *Hspg2* gains significance in light of a recent study comparing CAFs from genetically engineered mouse models of PDAC featuring mutant KRAS as well as mutant p53 (R172H) or p53 loss (43). The authors found that CAFs associated with p53-mutant PDAC are significantly more pro-metastatic and more effectively promote chemoresistance than CAFs from p53-null tumors, with both phenotypes driven by stromal expression of perlecan. In addition, the axon guidance cues expressed by PSC-derived CAFs include members of the Slit/Robo family, with potential implications for regulation of tumor innervation. We note that immune-modulatory cytokines and chemokines as well as genes that make up major histocompatibility complex class II (MHCII) are strongly enriched in the tdTomato^+^ CAF fraction, suggesting that the previously described inflammatory CAFs or iCAFs (28) and antigen-presenting CAFs or apCAFs (11) do not have a PSC origin.

**Figure 3.**
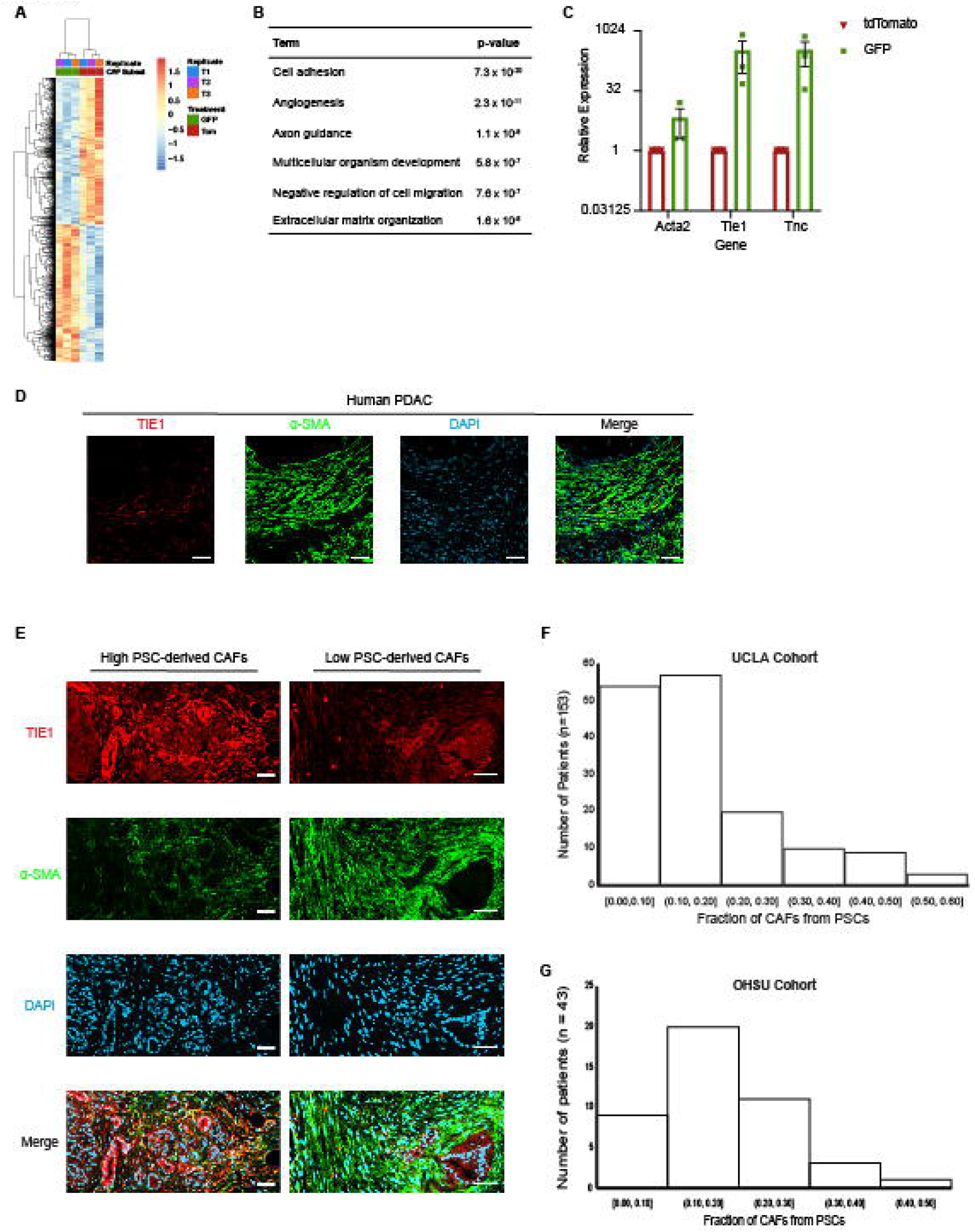
Mesenchymal lineage heterogeneity gives rise to transcriptional heterogeneity among PDAC CAFs. **(A)** Heatmap depicting differentially expressed genes in PSC-derived (GFP^+^) versus non-PSC-derived (tdTomato^+^) CAFs from KPC FC1199 PDAC in *Fabp4-Cre;Rosa26*^*mTmG*^ hosts (n = 3), identified by RNA-seq. **(B)** Gene ontology analysis identifying the top terms enriched in association with genes upregulated at least 2-fold in PSC-derived CAFs compared to non-PSC-derived CAFs. **(C)** qPCR for the indicated genes on PSC-derived and non-PSC-derived CAFs sorted from KPC FC1199 PDAC in *Fabp4-Cre;Rosa26*^*mTmG*^ hosts (n = 3) by FACS. Data were normalized to *36b4* and are presented as mean ± SEM. **(D)** Representative immunohistochemical staining of human PDAC for TIE1 and α-SMA (n = 5). Scale bar = 50 μm. **(E)** Representative images from a human PDAC microarray after immunohistochemical staining for TIE1 and α-SMA (n = 153). **(F)** Quantification of TIE1^+^α-SMA^+^ area out of total α-SMA^+^ area on each patient sample from the array. Different regions from the same patient were averaged together to yield one frequency per patient sample (4 punches per patient, 612 total tumor regions analyzed, 153 plotted here after averaging for each patient). **(G)** Quantification of TIE1^+^α-SMA^+^ area out of total α-SMA^+^ area using whole PDAC tissue sections (n = 43) from an independent patient cohort from that depicted in E & F.

We used our RNA-seq datasets to determine a marker combination for PSC-derived CAFs that could be used to analyze the frequency of this CAF population among PDAC patient samples, in hopes of validating the findings from our mouse models. For this, we selected the marker combination of α-SMA, a marker of the majority of CAFs and those in the myCAF population which include PSC-derived CAFs; and TIE1, which is highly expressed on endothelial cells but was unique to PSC-derived CAFs among total CAFs (Supplementary Fig. S2A). When we tested this marker combination on a small number of PDAC patient tumor sections, we noted a minor population of TIE1^+^α-SMA^+^ CAFs out of the expansive α-SMA^+^ population (Fig. 3D), consistent with patterns in our mouse models. To analyze the frequency of PSC-derived CAFs in human PDAC, including heterogeneity across a patient population, we obtained a PDAC tumor microarray containing four spatially distinct punches from each of 153 patient samples. We co-stained the array for TIE1 and α-SMA, and quantified the double-positive CAFs out of total α-SMA^+^ CAFs. We saw evidence of heterogeneity across this patient population, with some samples harboring relatively high levels of putative PSC-derived CAFs and others with very low levels (Fig. 3E). Quantification of CAF frequencies yielded two important conclusions (Fig. 3F): first, the vast majority of patient samples harbored putative PSC-derived CAFs at similar frequencies to those observed in our mouse models; and second, almost all of these patient samples had putative PSC-derived CAFs as the minority of CAFs in the tumor microenvironment, highlighting the extent of mesenchymal lineage heterogeneity. To validate these findings in an independent patient cohort, we obtained 43 PDAC tumor sections, here analyzing a lower “n” but whole sections instead of small punches on an array. Co-staining of these patient samples yielded similar results, including heterogeneity in putative PSC-derived CAF frequencies (Supplementary Fig. S2B), similar frequencies to those observed in mouse models in the majority of patient samples (Fig. 3G), and a minority of CAFs of a presumed PSC origin in all of these patient samples. Together, these results suggest that mesenchymal lineage heterogeneity underlies transcriptional heterogeneity among PDAC CAFs, and that this lineage heterogeneity is relevant to human PDAC.

### Targeted ablation of PSC-derived CAFs via retrograde ductal injection of viral Cre to unveil functional significance

We next wished to functionally interrogate PSC-derived CAFs to address whether their unique transcriptional profile translates to non-redundant roles in PDAC. To address this question, we aimed to ablate PSC-derived CAFs in established PDAC and analyze impacts on the tumor microenvironment. Crossing mice with a Cre-inducible diphtheria toxin receptor allele (*Rosa26*^*LSL-Hbegf/+*^, iDTR hereafter) (44) into our *Fabp4-Cre;Rosa26*^*mTmG*^ model together with diphtheria toxin (DT) treatment would lead to ablation of PSC-derived CAFs, but there were two obvious limitations of this model to overcome. First, as *Fabp4-Cre* mice feature high Cre activity in adipose tissue, this strategy would lead to systemic ablation of adipocytes and make our results difficult to interpret. Second, during characterization of our model, we noted a small population of CD45^+^GFP^+^ cells (about 2% of total intratumoral leukocytes, Supplementary Fig. S3A) which would also be targeted for ablation with this strategy. Notably, we did not observe CD45^+^GFP^+^ cells in normal pancreas of *Fabp4-Cre;Rosa26*^*mTmG*^ mice, suggesting that this Fabp4-expressing leukocyte population gets recruited into the tissue during tumor progression. To address these limitations and enable specific targeting of PSC-derived CAFs, we adapted published methods for retrograde ductal injection of viral particles directly into the pancreas (45, 46). This viral transduction approach achieves spatial control, limiting Fabp4-Cre to the pancreas and negating effects on adipose tissue, the hematopoietic system, and other potentially cell or tissue types with Fabp4-driven Cre activity; this also enables temporal control, introducing Fabp4-Cre in the adult mouse. To achieve this, we identified a minimal Fabp4 promoter and enhancer element that is highly active in primary PSCs but small enough to fit upstream of Cre within an AAV vector (see Methods). During optimization experiments, we found KP1 to be an optimal serotype to transduce PSCs *in vivo*, such that ductal injection of AAVKP1-Fabp4-Cre resulted in PSC labeling to a similar extent to that seen in mice harboring a *Fabp4-Cre* allele (Fig. 4A).

**Figure 4.**
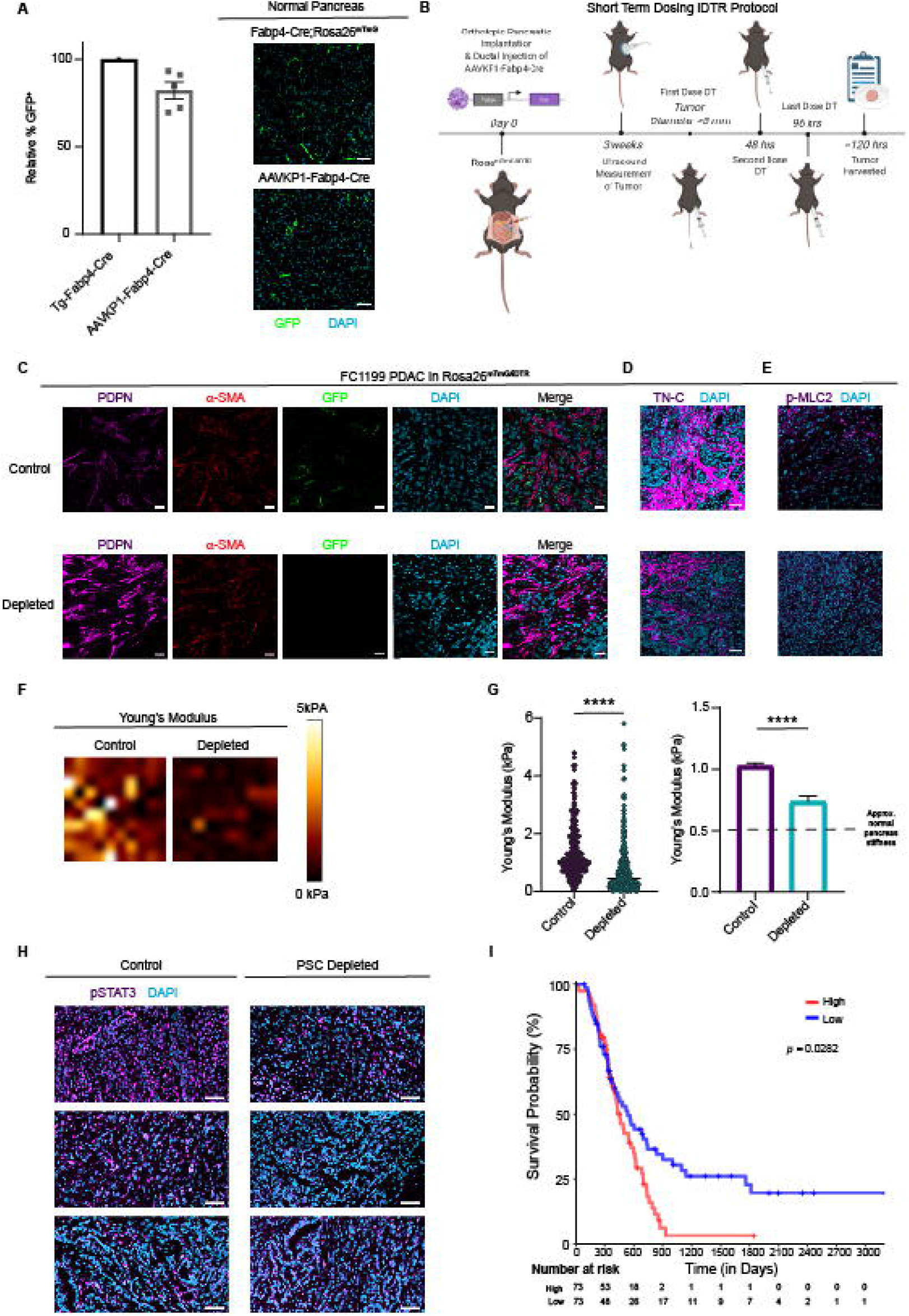
Targeted ablation reveals unique roles for PSC-derived CAFs in regulation of the extracellular matrix and mechanosignaling. **(A)** Immunohistochemical staining and quantification of GFP^+^ cells in normal pancreas tissue from *Fabp4-Cre;Rosa26*^*mTmG*^ mice and from *Rosa26*^*mTmG/iDTR*^ mice 7 days after intraductal injection with AAVKP1-Fabp4-Cre (n = 5). Data are presented as mean ± SEM. Scale bar = 50 μm. **(B)** Schematic of tumor modeling using intraductal injection of AAVKP1-Fabp4-Cre and orthotopic transplantation of KPC PDAC cells into *Rosa26*^*mTmG/iDTR*^ hosts. **(C)** Immunohistochemical staining for GFP, PDPN, and α-SMA of KPC FC1199 PDAC in *Rosa26*^*mTmG/iDTR*^ hosts with intraductal injection of AAVKP1-Fabp4-Cre, enrolled when tumors reached 5-6 mm in diameter and treated with PBS or DT for 5 days (n = 4). Scale bar = 20 μm. **(D)** Immunohistochemical staining for TNC of KPC FC1199 PDAC in AAVKP1-Fabp4-Cre-injected *Rosa26*^*mTmG/iDTR*^ hosts, enrolled at 5-6 mm in tumor diameter and treated with PBS or DT for 5 days (n = 3). Scale bar = 50 μm. **(E)** Immunohistochemical staining for p-MLC2 of PDAC samples as described in D. Scale bar = 50 μm. **(F)** Force maps generated by atomic force microscopy on KPC FC1199 PDAC in AAVKP1-Fabp4-Cre-injected *Rosa26*^*mTmG/iDTR*^ hosts (n = 3 per treatment group, control: 1063 data points, depleted: 717 data points), excised after 5 days of treatment with PBS or DT. **(G)** Quantification of Young’s modulus per AFM measurements on control and PSC-depleted PDAC as described in F. The dashed line on the graph on the right denotes the approximate stiffness of normal murine pancreas tissue. ****p < 0.0001 by unpaired t-test. **(H)** Immunohistochemical staining for p-STAT3 (Y705) of control and PSC-depleted PDAC harvested after 5 days of depletion (n = 3). Scale bar = 50 μm. **(I)** Kaplan-Meier plot depicting overall survival of PDAC patients with high versus low expression of a PSC-derived CAF ECM gene signature comprised of 99 genes (see Methods), plotting the upper versus lower quartile (n = 73 per arm).

We next moved to tumor modeling, for which we subjected *Rosa26*^*mTmG/iDTR*^ mice to a single surgery including ductal injection of AAVKP1-Fabp4-Cre and orthotopic injection of KPC PDAC cells (Fig. 4B). Established tumors in this model harbored the expected frequencies of GFP^+^ CAFs and lacked CD45^+^GFP^+^ cells (Supplementary Fig. S3B). To assess the functions of PSC-derived CAFs directly, as opposed to secondary impacts on other cells in the tumor microenvironment, we established tumors in our *Rosa26*^*mTmG/iDTR*^ hosts transduced with AAVKP1-Fabp4-Cre, enrolled when tumors reached 5-6 mm in diameter by high-resolution ultrasound, treated with vehicle or DT for just 5 days to acutely ablate PSC-derived CAFs, and harvested tumor tissue for analysis. Consistent with selective targeting of a numerically minor CAF population, the DT-treated tumors retained high levels of CAFs, including those expressing α-SMA (Fig. 4C). Based on our transcriptional profiling, we analyzed ECM components enriched in PSC-derived CAFs and found tenascin C (Fig. 4D, Supplementary Fig. S3C) significantly reduced upon PSC-derived CAF ablation. Total collagen abundance remained unchanged (Supplementary Fig. S3D), together suggesting that PSC-derived CAFs regulate specific components of the ECM. Consistent with a broader role in ECM regulation and mechanosignaling, phospho-myosin light chain 2 (p-MLC2) was markedly reduced upon ablation of PSC-derived CAFs (Fig. 4E). To test tumor stiffness directly, we analyzed control and PSC-depleted PDAC by atomic force microscopy, which revealed that tumor tissues were significantly softer upon ablation of the numerically minor PSC-derived CAF population (Fig. 4F,G; Young’s modulus for control = 1.02 ± 0.02 kPa, PSC-depleted = 0.74 ± 0.04 kPa; normal murine pancreas ≈ 0.50 kPa (47)). As tumor stiffness and tumor-promoting mechanosignaling have been shown to be facilitated in part by STAT3 in PDAC (18, 48-50), we analyzed levels of phospho-STAT3 (Y705) and found that these signaling events were markedly reduced upon PSC-derived CAF ablation (Fig. 4H). As these results suggested that PSC-derived CAFs drive the establishment of a tumor-promoting desmoplastic milieu, we developed an ECM signature specific to or enriched among PSC-derived CAFs per our RNA-seq datasets. While collagen and bulk tumor stromal density associated with a better prognosis in PDAC (25, 27), this PSC-associated ECM signature associated with a worse prognosis among PDAC patients (Fig. 4I, Supplementary Table S1). These results suggest that PSCs give rise to CAFs which regulate specific features of the stromal microenvironment associated with PDAC aggressiveness.

### Relationship between tumor genotype and stromal evolutionary routes with respect to CAF origins

Our results in PDAC patients suggested heterogeneity in PSC-derived CAF frequencies, and led us to question whether tumor genotype regulates stromal evolution from distinct cells of origin. In support of this notion, we observed significantly elevated *Hspg2* (perlecan) expression in PSC-derived CAFs compared to CAFs of other origins, raising the possibility that previously reported distinctions between CAFs associated with p53-mutant versus p53-null PDAC (43) reflect differential recruitment of PSCs into the CAF pool. To begin to test a relationship between cancer cell-intrinsic p53 status and stromal evolution, we compared two PDAC models from the same genetic background (C57BL/6J) and the same driver mutation in KRAS (G12D), but one featuring p53 R172H and the other featuring p53 loss. We harvested size-matched tumors from these models in *Fabp4-Cre;Rosa26*^*mTmG*^ hosts and found that total PDPN^+^ CAF frequencies were not different (Fig. 5A), but that PSCs made a significantly lower contribution to the CAF population in the context of p53-null PDAC (Fig. 5B,C). To assess a causal role for p53 status in regulation of stromal evolution, we generated isogenic models by using Cas9 and two different sgRNAs targeting *Trp53* to knock out p53 in a p53-mutant (R172H) parental line (Fig. 5D). We then transplanted these lines in pancreata of *Fabp4-Cre;Rosa26*^*mTmG*^ mice, harvested size-matched tumors, and analyzed CAF lineages by flow cytometry. We found that loss of p53 led to a reduction of PSC-derived CAF frequencies out of total PDPN^+^ CAFs (Fig. 5E) though we note that, for one of the p53-null lines, this trend did not reach statistical significance. Further, while p53-null tumors harbored a substantial CAF population as expected, p-MLC2 abundance was significantly reduced compared to p53-mutant tumors (Fig. 5F), consistent with a previous study (43) and with reduced PSC-derived CAF frequencies. These results suggest that cancer cell-derived factors stimulate stromal evolution, and that PDAC of distinct genotypes may be differentially responsive to stroma-targeted therapies independent of stromal density as a consequence of distinct mesenchymal cells of origin.

**Figure 5.**
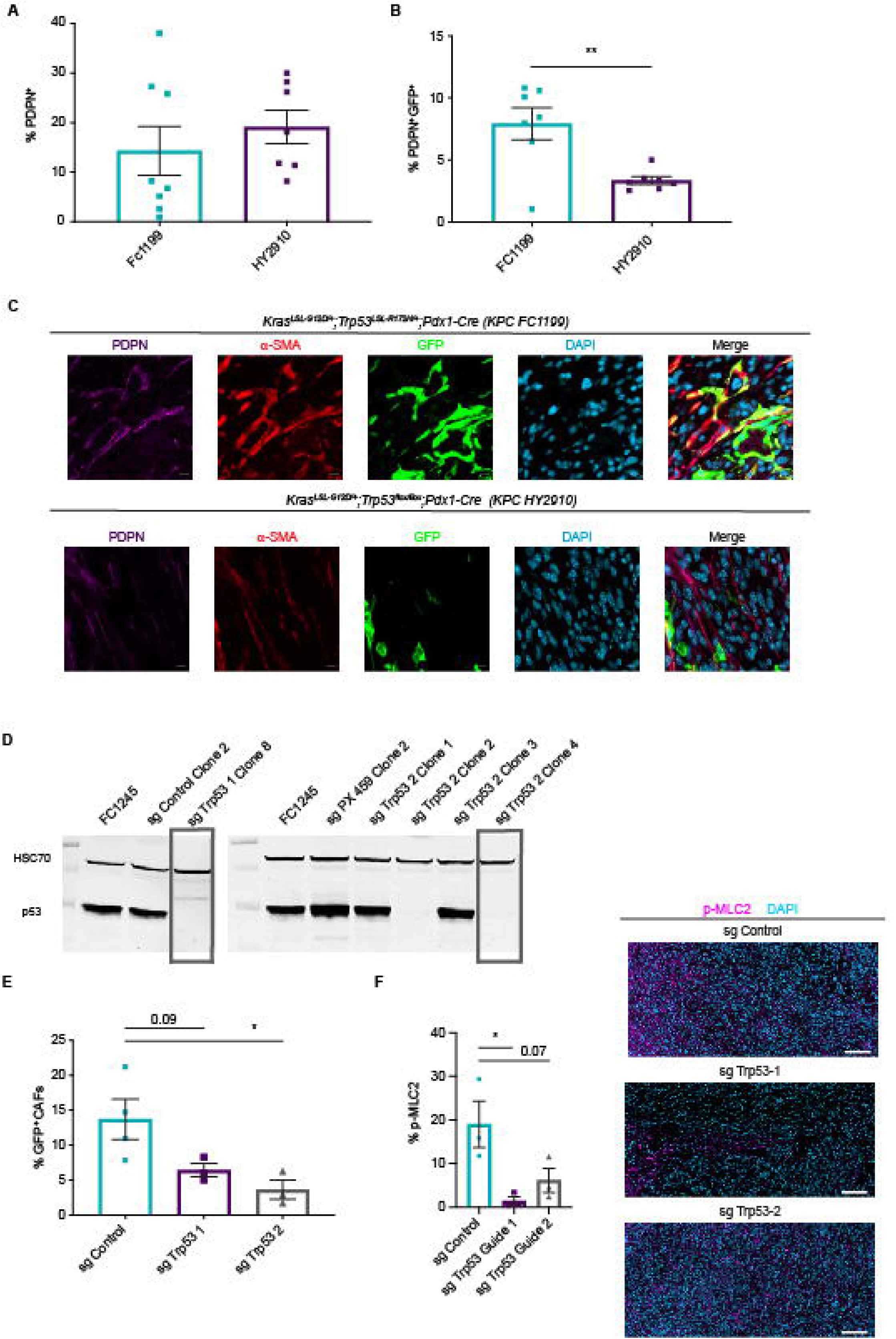
Tumor genotype with respect to p53 status influences stromal evolutionary routes. **(A)** Flow cytometry analysis of PDPN^+^ cells in size-matched KPC FC1199 (p53 R172H, n = 8) and HY2910 (p53-null, n = 7) PDAC in *Fabp4-Cre;Rosa26*^*mTmG*^ hosts. Data are presented as mean ± SEM. **(B)** Flow cytometry analysis of PDPN, GFP, and tdTomato in the tumors described in A to quantify the percent of CAFs derived from PSCs. Data are presented as mean ± SEM. **(C)** Immunohistochemical staining for GFP and PDPN on KPC FC1199 and HY2910 PDAC in *Fabp4-Cre;Rosa26*^*mTmG*^ hosts (n = 3). Scale bar = 10 μm. **(D)** Western blots for p53 and HSC70 (loading control) using whole cell lysates from parental KPC FC1245 (p53 R172H) cells or derivative lines transfected with control plasmid or 1 of 2 sgTrp53 sequences. **(E)** Flow cytometry analysis of PDPN, GFP, and tdTomato in size-matched control (n = 4) and sgTrp53 (n = 3 per line) PDAC in *Fabp4-Cre;Rosa26*^*mTmG*^ hosts. Data are presented as mean ± SEM. *p < 0.05 by one-way ANOVA. **(F)** Immunohistochemical staining for p-MLC2 in size-matched control and sgTrp53 PDAC in *Fabp4-Cre;Rosa26*^*mTmG*^ hosts (n = 3). Scale bar = 100 μm. *p < 0.05 by one-way ANOVA.

## Discussion

The discovery of stellate cells in the pancreas in 1998 provided a crucial basis for our understanding of the cellular source(s) of pancreatic fibrosis (39). As hepatic stellate cells serve as the dominant cellular source of fibrosis in the context of liver injury (41) and PSCs were shown to have similar fibrogenic potential to their counterparts in the liver (32), it was reasonable to speculate that PSCs are the cellular source of pancreatic fibrosis and of the extensive desmoplastic reaction in PDAC (31). Indeed, activation of PSCs in culture leads to induction of a transcriptional program consistent with CAF features (36) and, more recently, PSCs were shown to give rise to the previously defined iCAF and myCAF subtypes in culture upon exposure to defined soluble cues and growth substrates (35). In light of the plasticity of this cell type and their presumed role as dominant contributors to the PDAC stroma, we expected our genetic means to track and ablate PSCs and derivative CAFs would target most or all CAFs in the PDAC microenvironment. We were surprised to find that PSCs only give rise to a small minority of CAFs in PDAC, with frequency dependent in part on tumor genotype. As we come to further understand the functions of this CAF population, we will further analyze clinical cohorts, leveraging heterogeneity in PSC-derived CAF frequency among patient samples to investigate potential stratification strategies or tailored therapies informed by CAF cell of origin.

As PSC-derived CAFs have a transcriptional profile distinct from CAFs of other origins, our results in the context of the recent literature suggest that CAF transcriptional heterogeneity results from (at least) two sources: signaling gradients differentially regulating common cells of origin, and mesenchymal lineage heterogeneity. The results of our PSC-derived CAF ablation experiments suggest that functional heterogeneity underlies transcriptional heterogeneity. While transcriptional and phenotypic plasticity among CAFs likely poses some limitations to the feasibility of targeting specific subsets therapeutically, our model enabling targeted ablation of a defined CAF subset raises the possibility that CAF subsets have sufficient functional distinctions that targeting these subsets in preclinical models, to understand their role in PDAC biology, and differentially targeting them therapeutically may indeed be possible. This notion is supported by a recent study employing the potent smoothened antagonist LDE225 to effectively inhibit Hedgehog signaling in mouse models of PDAC (20). This study showed that myCAFs are partially dependent on the Hedgehog pathway, such that LDE225 treatment markedly skewed that PDAC CAF population to reduce myCAFs and increase iCAFs. Impacts of LDE225 treatment on tumor-infiltrating T cells suggest potential immune-suppressive functions of iCAFs and, as the unique functions of these distinct CAF subsets further come to light, targeting specific CAFs may emerge as viable combination therapeutic strategies. To this end, it will be important to extend the functional analyses performed here to additional CAF populations, pending development of relevant models, to better understand their distinct roles in the tumor microenvironment and inform on potential targets for therapeutic intervention.

The high α-SMA expression and ECM production characteristic of PSC-derived CAFs suggest that they fall within the previously described myCAF (11, 28) or TGFβ CAF (12) population. However, as total collagen abundance remains unchanged and α-SMA^+^ CAFs remain highly abundant upon PSC-derived CAF ablation, this numerically minor populations seems to be but a subset of the broader myCAF CAF pool. As PSC-derived CAFs regulate specific ECM components and biophysical properties implicated in tumor aggressiveness and metastatic progression (18, 43), we speculate that PSCs give rise to a tumor-promoting myCAF subset, to be investigated in depth in subsequent studies. However, in light of the apparent dependency of myCAFs on Hedgehog signaling and the detrimental effects of long-term genetic or pharmacologic SHH inhibition (23, 24), it seems likely that another subset of myCAFs is tumor-suppressive and promotes a more differentiated and less aggressive PDAC phenotype. The tumor-suppressive potential of other myCAF constituents is consistent with prior studies providing correlative evidence for heightened immune suppression upon reduction in overall myCAF frequency in the PDAC microenvironment (13, 20). These tumor-suppressive CAF populations and their homeostatic or beneficial functions will be important to identify, as maintaining these functions should be a goal of future stroma-targeted therapies.

As our data define PSCs as numerically minor contributes to the CAF population, an important question to address is the cellular origin of the majority of PDAC CAFs. Analysis of single-cell RNA-seq data from murine PDAC together with publicly available transcriptional profiles suggests that the previously described apCAFs in fact derive from mesothelial cells (12). As for the remaining CAFs, including iCAFs and the non-PSC-derived myCAFs, the origins remain unclear but may come from pancreas-resident fibroblast populations or potentially from the bone marrow. While bone marrow-derived mesenchymal stromal cells are important contributors to the CAF population in other cancer types, including breast cancer (51), the contribution of bone marrow progenitors to PDAC CAFs remains to be determined. Importantly, a recent study employed lineage tracing and identified two distinct fibroblast populations in normal pancreas tissue, one marked by the Hedgehog-responsive transcription factor Gli1 and the other marked by the tissue-restricted mesenchymal transcription factor Hoxb6 (52). These factors label distinct fibroblast populations in the pancreas which are similar in frequency, but in the early stages of pancreatic carcinogenesis, the Gli1^+^ fibroblast population expands considerably while the Hoxb6^+^ population does not, such that the pancreas-resident, Gli1^+^ fibroblasts are major contributors to the stroma associated with precursor lesions. Future studies to define the mechanisms shaping PDAC CAF heterogeneity and the unique functions of distinct CAF subsets will improve our understanding of how the tumor microenvironment impacts PDAC progression, and perhaps point to important stromal targets for therapeutic intervention.

## Methods

### Animals

All experiments performed in mice were reviewed and overseen by the institutional animal care and use committee at Oregon Health & Science University in accordance with NIH guidelines for the humane treatment of animals. *Fabp4-Cre* (005069) and *Rosa26*^*mTmG*^ (007676) mice from Jackson Laboratory were used for PSC analyses and orthotopic transplantation experiments at 8-12 weeks of age, including male and female mice. *Rosa26*^*mTmG*^ and iDTR (007900) mice from Jackson Laboratory were used for retrograde ductal injection and orthotopic transplantation for PSC ablation experiments at 8-12 weeks of age, including male and female mice.

### Human tissue samples

Human patient PDAC tissue samples donated to the Oregon Pancreas Tissue Registry program (OPTR) with informed written patient consent (IRB approved, IRB00003609) in accordance with full ethical approval by the Oregon Health & Science University Institutional Review Board were provided by the OHSU Brenden-Colson Center for Pancreatic Care and the Knight BioLibrary, upon pathology review by Dr. Rosemary Makar and secondary review by Dr. Christopher Corless.

The PDAC tumor microarray was described previously (53). Human PDAC specimens were obtained from patients who underwent surgical resection of primary PDAC, under IRB-approved protocol IRB11-000512. This study was conducted under strict compliance with institutional ethical regulations. The study had minimal risk per the IRB protocol and thus informed consent was not necessary.

### Immunohistochemistry

For mouse tissue harvest, mice were anesthetized and euthanized according to institutional guidelines. Pancreas tissue or tumors were excised carefully and fixed overnight in 10% neutral buffered formalin, or embedded in OCT and frozen at −80°C. Fixed tissues were paraffin-embedded, sectioned, deparaffinized and rehydrated through an ethanol series and ultimately in PBS. Following antigen retrieval, tissue samples were blocked for two hours at room temperature in blocking solution (8% BSA) and transferred to a carrier solution (1% BSA) containing diluted antibodies. Sections were incubated overnight at room temperature and then washed five times for 5 minutes each in PBS. Secondary Alexa-fluor conjugated antibodies diluted in the same carrier solution (1:200) were added to the sections for two hours at room temperature. Sections were then washed five times for five minutes each in PBS, autofluorescence quenched with the TrueVIEW reagent (Vector Laboratories), stained with DAPI, and mounted with Vectashield mounting medium. Fresh-frozen tissues were sectioned, fixed, and used to stain for CD45 in FC1199 tumors and for NG2 in normal pancreas tissue, then counterstained with DAPI-containing Vectashield mounting medium for fluorescence microscopy. Antibodies used for immunohistochemistry were as follows: CD31: Abcam ab28364, GFP: Cell Signaling Technology 4B10 (mouse) or D5.1 (rabbit), CD45: Abcam ab25386, NG2: EMD Millipore AB5320, α-SMA: Thermo Fisher MA511547, PDPN: Thermo Fisher 14-5381-81, TIE1: Abcam ab111547 (mouse tissues), TIE1: Thermo Fisher PA527903 (human tissues), TNC: Abcam ab108930, pMLC2: Cell Signaling Technology 3674, pSTAT3: Cell Signaling Technology 9145. Stained tissues were imaged using a laser-scanning confocal inverted microscope (LSM 880, Carl Zeiss, Inc.) and a 40x/1.1 NA water objective or 63x/1.4 NA oil objective was used to image the samples. Slides were scanned using a Zeiss Axio Scan.Z1 and quantified using QuPath or Aperio software.

For quantification of TIE1/α-SMA colocalization on human PDAC sections, images were acquired on an AxioScan.Z1 using a 10x 0.45 NA plan-apochromat lens. Fluorochromes were excited with a Colibri 7 light source (Carl Zeiss), and excitation and emission light was passed through the following Zeiss filter sets for the appropriate channel: DAPI-96 HE; Alexa-fluor 488-HE 38; Alexa-fluor 594-71 HcRed. Images were analyzed for colocalization in ZEN v2.3 (Carl Zeiss). Thresholds for the AF594 and AF488 channels were set by eye for each slide (three slides total) for the TMA analysis. Within each slide, the same thresholds were used across all tissues. Dynamic range was set to a 14-bit image (16384 maximum intensity).

### Fluorescence activated cell sorting and flow cytometry

For flow cytometry analysis of normal pancreas tissue, pancreata were harvested, briefly minced with scissors, and digested with 0.02% Pronase (Sigma-Aldrich), 0.05% Collagenase P (Sigma-Aldrich), and 0.1% DNase I (Sigma-Aldrich) in Gey’s balanced salt solution (GBSS, Sigma-Aldrich) at 37°C for 20 mins. After dissociation, tissue was triturated until large pieces were no longer visible, and the resulting cell suspension was filtered through a 100-μm nylon mesh. Cells were then washed with GBSS, pelleted, subject to red blood cell lysis in ACK lysis buffer (Thermo Fisher) for 3 mins at room temperature, washed in FACS buffer (PBS containing 2% FBS), pelleted, and resuspended in FACS buffer for flow cytometry to analyze vitamin A positivity based on autofluorescence using the 405 nm laser on a BD Fortessa flow cytometer. To analyze endothelial cells, following red blood cell lysis, cells were incubated with CD16/CD32 antibody (BD Biosciences 553141) to block Fc receptors for 2 mins at room temperature, then stained with a PerCP/Cy5.5-conjugated CD31 antibody (BioLegend 102522) for 30 mins on ice. Stained cells were washed with cold FACS buffer, pelleted, and resuspended in cold FACS buffer for flow cytometry analysis.

For analytical flow cytometry or FACS on PDAC tissues, tumors were harvested, minced with scissors, and dissociated in DMEM containing 1 mg/ml Collagenase IV (Thermo Fisher), 0.1% Soybean Trypsin Inhibitor (Thermo Fisher), 50 U/ml DNase I (Sigma-Aldrich), and 0.125 mg/ml Dispase II (Thermo Fisher) at 37°C for 1 hr. Digested tissues were pelleted, resuspended in 0.25% Trypsin in DMEM, and incubated at 37°C for 10 mins, then washed with cold DMEM containing 10% FBS and pelleted. Resuspended cells were filtered through a 100 μm cell strainer, pelleted, washed with DMEM + 10% FBS, and pelleted again. Cells were resuspended in ACK lysis buffer as described above, washed with FACS buffer, pelleted, and resuspended in FACS buffer. Fc block was performed as described above, then cells were stained with antibodies for 30 mins on ice: biotinylated anti-PDPN (BioLegend 127404), biotinylated anti-PDGFRα (anti-CD140a, BioLegend 135910), PerCP/Cy5.5-conjugated anti-CD31 (BioLegend 102522), and/or PE/Cy7-conjugated anti-CD45 (BioLegend 103113). Cells were then washed with FACS buffer and pelleted. When biotinylated antibodies were used, incubated with APC Streptavidin (BD 554067) for 30 mins on ice, washed with FACS buffer, pelleted, and resuspended in FACS buffer. After staining for the purpose of CAF sorting and subsequent RNA-seq, RNase inhibitor (New Englad Biolabs M0314, 1:40) was added to the cell suspension, and CAFs were sorted into TRIzol LS (Thermo Fisher). Flow cytometry analysis was performed on a BD Fortessa, while CAF sorting was performed on a BD FACSAria Fusion. Flow cytometry data were analyzed with FlowJo software. GFP^+^ or tdTomato^+^ CAF frequencies were calculated out of the total GFP^+^ plus tdTomato^+^ population, excluding any cells lacking either lineage label which represent tumor cells.

### Orthotopic PDAC experiments

Male and female mice at 8-12 weeks of age were used as hosts for PDAC orthotopic transplantation, using genotypes described in the Results section and figure legends. Mice were anesthetized with ketamine and xylazine, and pancreata were injected with 5 × 10^3^ FC1199 or FC1245 cells (lines provided by Dr. David Tuveson), 1 × 10^5^ 4662 cells (provided by Dr. Robert Vonderheide), or 1 × 10^4^ HY2910 cells (provided by Dr. Haoqiang Ying), all derived from primary PDAC in *Kras*^*LSL-G12D/+*^*;Trp53*^*LSL-R172H/+*^*;Pdx1-Cre* mice (FC1199, FC1245, 4662) or *Kras*^*LSL-G12D/+*^*;Trp53*^*flox/flox*^*;Pdx1-Cre* mice (HY2910) of a C57BL/6J genetic background. Orthotopic transplantation was performed as previously described (54). For PSC-derived CAF ablation experiments, pancreata were imaged beginning 14 days after transplantation by high-resolution ultrasound using the Vevo 770 imaging system; mice were enrolled on study when tumors reached 5-6 mm in diameter. Enrolled mice were treated with sterile PBS or 25 ng/g diphtheria toxin (List Biological Laboratories) by intraperitoneal injection every 2 days and tumors were harvested on day 5 post-enrollment for analysis.

### Retrograde ductal AAV delivery

A promoter and enhancer element upstream of mouse *Fabp4* (from Addgene #8858) was cloned into pAAV-iCre-WPRE (Vector Biosystems) upstream of Cre. KP1-serotyped AAV-Fabp4-Cre was generated and tittered by the OHSU Molecular Virology Core. AAVKP1-Fabp4-Cre viral stock was diluted to 1 × 10^10^ viral genomes/ml in 10 μg/ml DEAE-Dextran (Sigma-Aldrich) in PBS, incubated for 30 mins at room temperature, then placed on ice. Retrograde ductal injections were performed as previously described (45). Mice were anesthetized with ketamine and xylazine until non-responsive to toe pinch. Feet were taped down with surgical tape on a sterile mat. Entire abdomen was shaved, and the shaved area cleaned with 70% ethanol and betadine. A midline incision was made about 1 inch long, and incision edges were secured with hemostatic forceps. Non-weave sterile gauze was moistened with sterile PBS. Intestines were gently extracted using circle forceps, and gently laid onto damp gauze. Intestines were slowly extracted until duct was visible. Exposed intestines were covered with damp gauze to keep moist, and sterile PBS was continually added throughout the procedure as necessary to keep gauze and tissue from drying out.

One microvascular clip was placed on the cystic duct near the gallbladder. The sphincter of Oddi was located; a 30G insulin syringe was loaded with 100 μl viral solution (to inject 1 × 10^9^ viral genomes per mouse), and the needle inserted through the sphincter of Oddi into the common bile duct up to its convergence with the cystic duct, about halfway to the clip. Viral solution was slowly injected over the course of 2 mins. The needle was left in place for 30 sec after completion of the injection, then slowly and gently removed. The clip was then removed. Intestines were carefully returned to the abdomen, muscle layer closed with vicryl sutures, and skin closed with sterile suture clips. When performed together with orthotopic transplantation of PDAC cells, cells were injected after removal of the syringe and clip.

### Atomic force microscopy

Fresh tumor tissues were placed in the middle of a disposable cryomold and embedded in Optical Cutting Temperature (OCT) compound. Tissues were sectioned with a cryostat at a thickness of 50 μm and adhered to positively charged microscope slides. Prior to measurements, samples were thawed at room temperature in DMEM/F-12 supplemented with 10% FBS and a protease inhibitor cocktail (Sigma Aldrich 11836170001). Atomic force microscopy (AFM) measurements were performed with a NanoWizard 4 XP BioScience with HybridStage (Bruker) mounted on a Zeiss Axio Observer inverted optical microscope. Dulled triangular silicon nitride ScanAsyst-Fluid probes were used (κ = 0.7 N/m; Bruker) at a maximum applied force of 4 nN and approach speed of 5 μm/s. Each probe was calibrated with the thermal oscillation method prior to measurements on each tissue. Force curves were analyzed to obtain the Young’s modulus using the JPK Data Processing Software package with the Hertz/Sneddon model assuming an incompressible tissue and a Poisson’s ratio of 0.5 with tip parameters provided by the manufacturer’s specifications. Three independent samples were imaged from each treatment group at three locations at varying tissue depths with a scan size of 100 μm^2^ with force measurements spaced every 10 μm.

### Survival analysis of a PSC ECM gene signature

The PSC-derived CAF ECM signature genes (Supplementary Table S1) were defined as the overlap of the genes annotated in the Reactome Extracellular Matrix Organization pathway and the upregulated genes in GFP^+^ CAFs (FDR < 0.01 and fold change > 1.25) per our RNA-seq analysis. In the signature survival analysis, 297 samples were collected with survival time from the OHSU Brenden-Colson Center for Pancreatic Care Tempus dataset. The gene set variation analysis (GSVA) algorithm with the default settings, as implemented in the GSVA R package (version 1.34.0), was applied to calculate the gene signature score for each sample. Next, the samples were stratified into two groups based on the quantile values of the signature scores (upper quartile versus lower quartile). Survival curves of these two groups of patients were analyzed by Kaplan-Meier method with statistical significance calculated using the log-rank test. The Kaplan-Meier estimator and log-rank test were calculated in the survival R package (version 3.2-3).

### Statistical analysis

Statistical analyses were performed using GraphPad PRISM software. Student *t* test was used to compare two groups to each other. One-way ANOVA was performed when multiple conditions were compared for one variable. Tukey *post hoc* tests were used after ANOVA analyses to perform multiple group comparison. Analysis with a p-value < 0.05 was considered statistically significant.

## Data availability

All sequence data from this study have been deposited in the publicly available Gene Expression Omnibus under accession number GSE143805.

## Acknowledgements

We thank all members of the Sherman lab, Jae Myoung Suh, and Sara Courtneidge for helpful conceptual input on this work; and Markus Grompe, Hiroyuki Nakai, Shin-Heng Chiou, and Monte Winslow for guidance on the intraductal AAV delivery approach. This work was supported by the OHSU Molecular Virology Core, Knight BioLibrary (and particularly by operations supervisor Danielle Galipeau and by Christopher Corless in his capacity as Chief Medical Officer of the Knight Diagnostic Laboratories), Flow Cytometry Shared Resource, Massively Parallel Sequencing Shared Resource, Advanced Light Microscopy Shared Resource, and Histopathology Shared Resource, with core facility support from the Knight Cancer Institute Cancer Center Support Grant P30 CA069533. Funding to support this study came from NIH grant T32 GM071338 (to E.H.), and NIH grant R01 CA250917, DOD Peer Reviewed Cancer Research Program grant W81XWH-18-1-0437, and a Pew-Stewart Scholar Award (all to M.H.S.).

## Author contributions

E.H. and M.H.S. conceived the project. E.H. and M.B. managed mouse breeding for the study, and performed the in vivo experiments. E.H., R.C.C., M.K.O., C.O., S.B., J.M.F., A.S., and M.H.S. performed immunohistochemical staining and/or analyzed stained tissues. C.C.D. performed atomic force microscopy, and analyzed the results and generated data together with S.R.H. E.H. and C.O. performed cell culture and molecular biology experiments. W.H. analyzed the RNA-seq data. R.M. provided pathology assessment of human PDAC samples. D.W.D. provided the human PDAC microarray. D.S. and Z.X. generated a gene signature from our RNA-seq data and analyzed prognostic value using human PDAC RNA-seq data. E.H. and M.H.S. wrote the manuscript with input from all authors.

## Figure legends

**Figure S1, related to Figure 1.**

**(A)** Flow cytometry analysis of CD31, GFP, and tdTomato on normal pancreas tissue from *Fabp4-Cre;Rosa26*^*mTmG*^ mice (n = 3). Data are presented as mean ± SEM. **(B)** Representative flow cytometry plot showing GFP^+^ and tdTomato^+^ populations in normal pancreas from *Fabp4-Cre;Rosa26*^*mTmG*^ mice used for FACS. **(C)** Immunofluorescence staining for Desmin and GFP on PSCs isolated from *Fabp4-Cre;Rosa26*^*mTmG*^ mice by density centrifugation. Scale bar = 10 μm. **(D)** Representative flow cytometry plot showing vitamin A^+^ cells in normal pancreas from *Fabp4-Cre;Rosa26*^*mTmG*^ mice.

**Figure S2, related to Figure 3.**

**(A)** Immunohistochemical staining for TIE1 and α-SMA in KPC FC1199 PDAC in *Fabp4-Cre;Rosa26*^*mTmG*^ mice (n = 3). Scale bar = 50 μm. **(B)** Representative immunohistochemical staining of human PDAC tissue sections (n = 43) for TIE1 and α-SMA. Scale bar = 100 μm.

**Figure S3, related to Figure 4.**

**(A)** Flow cytometry analysis of CD45 and GFP in KPC FC1199 PDAC in *Fabp4-Cre;Rosa26*^*mTmG*^ mice (n = 4). Data are presented as mean ± SEM. **(B)** Flow cytometry analysis of CD45, GFP, and tdTomato in KPC FC1199 PDAC in *Rosa26*^*mTmG/iDTR*^ mice transduced with intraductal AAVKP1-Fabp4-Cre (n = 3). Data are presented as mean ± SEM. **(C)** Immunohistochemical staining for TNC in KPC FC1245 PDAC in *Rosa26*^*mTmG/iDTR*^ mice transduced with intraductal AAVKP1-Fabp4-Cre and treated with PBS or DT for 5 days (n = 3). Scale bar = 10 μm. Data are presented as mean ± SEM. *p < 0.05 by unpaired t-test. **(D)** Trichrome staining and quantification of aniline blue signal (collagens) of KPC FC1199 PDAC in AAVKP1-Fabp4-Cre-injected *Rosa26*^*mTmG/iDTR*^ hosts, enrolled at 5-6 mm in tumor diameter and treated with PBS or DT for 5 days (n = 5). Scale bar = 50 μm.

