## Supplementary Methods for "Mesenchymal Lineage Heterogeneity Underlies Non-Redundant Functions of Pancreatic Cancer-Associated Fibroblasts"

Helms et al.

### Supplementary Methods

#### PSC isolation and analysis

PSCs were isolated from healthy pancreas tissue from 8 week old *Fabp4-Cre;Rosa26*$^{mTmG}$ mice as previously described (1). Upon retrieval from the density gradient interface, PSCs were plated on glass coverslips in DMEM containing 10% fetal bovine serum. On day 2 of culture, cells were fixed in 4% paraformaldehyde for 15 mins at room temperature, washed three times with PBS, and permeabilized with 0.1% Triton X-100 for 10 mins at room temperature. Following permeabilization, coverslips were blocked for 1 hr at room temperature in blocking solution (8% BSA), then transferred to a carrier solution (1% BSA) containing diluted antibodies against Desmin and GFP (Desmin: Cell Signaling Technology D93F5, GFP: Cell Signaling Technology 4B10). Coverslips were incubated with the primary antibody solution for 3 hr at room temperature, then washed five times for 5 mins each in PBS. Secondary Alex-fluor conjugated secondary antibodies were diluted in the same carrier solution (1:400) and added to the coverslips for 1 hr at room temperature. Coverslips were then washed five times for 5 mins each in PBS and mounted with Vectashield mounting medium containing DAPI. Cells were imaged using the LSM 880 confocal microscope described above, and co-stained cells out of total Desmin-positive cells were scored manually.

#### Gene expression analysis by qPCR

Total RNA was isolated using TRIzol reagent (Thermo Fisher) per manufacturer's instructions. The isolated total RNA (1 μg) was reverse-transcribed to produce cDNA using iScript Reverse Transcription Supermix kit (Bio-Rad). Real-time PCR was performed using SYBR Green supermix (Bio-Rad). The cDNA sequences for genes of interest were obtained from the mouse genome assembly (http://genome.ucsc.edu) and gene-specific primers were designed using the Primer3 program (http://frodo.wi.mit.edu/primer3/primer3_code.html). Relative gene expression was normalized using the 36B4 housekeeping gene. The following primer sequences were used: 36B4 (*Rplp0*): F: 5'-GTGCTGATGGGCAAGAAC-3', R: 5'-AGGTCCTCCTTGGTGAAC-3'; *Ins1*: F: 5'- TATAAAGCTGGTGGGCATCC-3', R: 5'-GGGACCACAAAGATGCTGTT-3'; *Fabp4*: F: 5'- GATGGTGACAAGCTGGTGGT-3', R: 5'- AATTTCCATCCAGGCCTCTT-3'; *Des*: F: 5'- ACTTGACTCAGGCAGCCAAT-3', R: 5'- ATCCTCCAGCTCCCTCATCT-3'; *Krt19*: F: 5'- TGACCTGGAGATGCAGATTG-3', R: 5'- AATCCACCTCCACACTGACC-3'; *Vim*: F: 5'- ACGGTTGAGACCAGAGATGG-3', R: 5'- TCTTGCGCTCCTGAAAAACT-3'; *Cspg4*: F: 5'- TTACCTTGGCCTTGTTGGTC-3', R: 5'- GAGCTGGAGCAAGAGATGGT-3'; *Prss3*: F: 5'- TCTGTCCCCTACCAGGTGTC-3', R: 5'- GTTGGGGTGCTTGATGATCT-3'; *Tie1*: F: 5'- TGCCCTTTTAGCCTTGGTGT-3', R: 5'- AGGATGGTCTCTTCACCCGA-3', *Tnc*: F: 5'- CCTGGACGGCATCGGAGAAT-3', R: 5'- TTGTTTGGTGCCCTTGAGTGA-3'; *Acta2*: F: 5'-TCAAGGAGAAGCTGTGCTATGT-3', R: 5'- TTCGTGGATGCCCGCTGA-3'

**RNA-seq**

RNA was isolated from sorted GFP$^+$ and tdTomato$^+$ CAFs (n = 3) using TRIzol and the Qiagen RNeasy Mini Kit with on-column DNase digestion for clean-up. Libraries were created using the SMART-Seq v4 Ultra Low Input RNA Kit for Sequencing (Takara Bio USA, Inc.). The input was 10 ng and 8 PCR cycles were used for the prep. Libraries were quantified using KAPA qPCR and pooled in an equimolar ratio. Libraries were sequences on the Illumina HiSeq 2500 on a Single Read Flow Cell using Illumina HiSeq SR cBot Cluster Kit v4 and HiSeq SBS v4 reagents.

RNA-seq reads were aligned to the human reference genome (GRCh38, release 84) using STAR (version 2.5.2b) with default parameters (2). The STAR “GeneCounts” module was used to quantify the number of reads mapping to each gene. Gene expression quantified by read counts from STAR were used as input into DESeq2 for differential expression gene (DEG) analysis (3). Subsequent DEG analyses were performed using the 12,749 genes with counts per million (cpm) greater than 1 in at least four samples. The DEGs were called with a false discovery rate (FDR) less than 0.05. In the DESeq2 package, counts were normalized using the variance stabilizing transformation (VST) module in DESeq2 for downstream analyses. For heatmap generation, differentially expressed genes were clustered using Pearson correlation distance and the complete clustering method from the pheatmap R package.

**Cell culture**

FC1199 and FC1245 (provided by Dr. David Tuveson), 4662 (provided by Dr. Robert Vonderheide), and HY2910 (provided by Dr. Haoqiang Ying) PDAC cell lines were derived from primary murine PDAC as described above. Cell lines were routinely passaged in DMEM (Thermo Fisher) containing 10% FBS (HyClone) for no more than 25-30 passages. Primary PSCs were also cultured in DMEM containing 10% FBS after isolation from mouse pancreas and until fixation and analysis. Cell lines were routinely tested for *Mycoplasma* at least monthly (MycoAlert Detection Kit, Lonza). Cell line authentication was not performed.

**Generation of p53-null PDAC cell lines**

The pSpCas9(BB)-2A-Puro(PX459) v2.0 plasmid (Addgene #62988) was used to clone guide sequences targeting *Trp53* per supplier's protocol: sgRNA #1: CACCGACCCTGTCACCGAGACCCC, sgRNA #2: CACCGGGAGCTCCTGACACTCGGA. The FC1245 PDAC cell line was transfected with control plasmid or plasmid containing either of the sg*Trp53* sequences and subject to selection with 2 μg/ml puromycin for 4 days. Single-cell clones were expanded and screened for p53 expression by Western blot, and clones with loss of p53 protein were selected for *in vivo* experiments.

**Western blotting**

For all Western blots, cells were harvested in ice-cold PBS and lysed in RIPA buffer containing phosphatase and protease inhibitors. Proteins were resolved on a 4-12% Bis-Tris NuPAGE gel (Invitrogen), then transferred onto a PVDF membrane. Membranes were probed with primary antibodies and infrared secondary antibodies: p53 (Cell Signaling Technology 2524), HSC70 (Santa Cruz sc-7298), anti-rabbit Alexa Fluor Plus 680 (Thermo Fisher A32734) and anti-mouse Alexa Fluor Plus 800 (Invitrogen A32730).

Protein bands were detected using the Odyssey CLx infrared imaging system (LICOR Biosciences).
