## Supplementary figures and images for "Mesenchymal Lineage Heterogeneity Underlies Non-Redundant Functions of Pancreatic Cancer-Associated Fibroblasts"

### Supplementary Figure 1

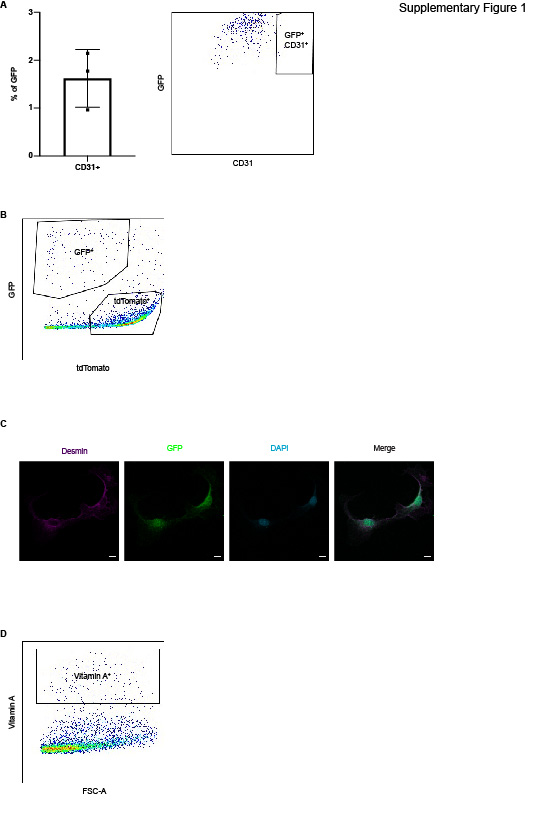

### Supplementary Figure 2

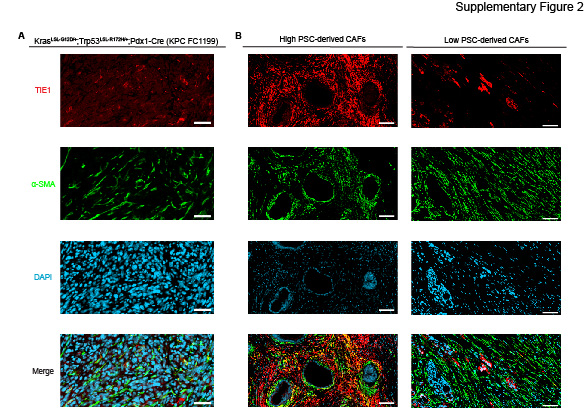

### Supplementary Figure 3

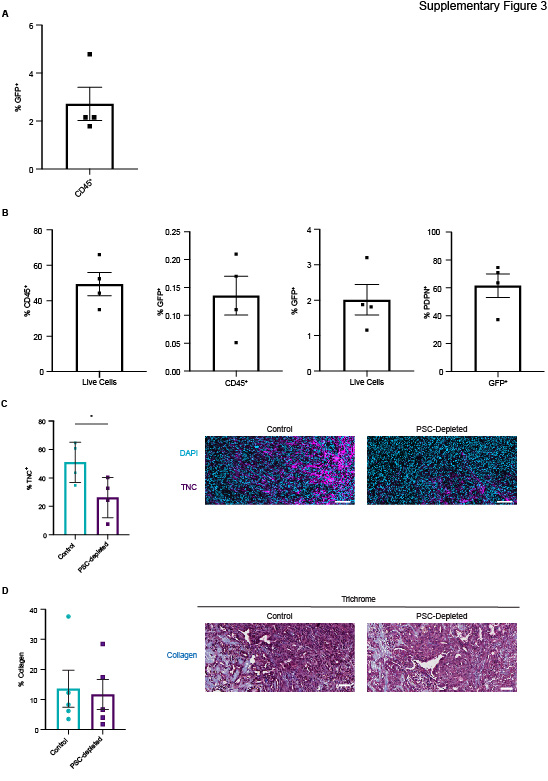
